# Cryo-EM complex structure of active GPR75 with a nanobody

**DOI:** 10.1101/2022.08.18.503988

**Authors:** Zilin Lv, Yuntong He, Yuning Xiang, Jing Li, Shuhao Zhang, Fanhao Meng, Baoliang Lan, Hanbo Guo, Dong He, Yanxia Wang, Huimin Zhao, Wei Zhuo, Yujie Liu, Xiangyu Liu, Xiaodan Ni, Jie Heng

## Abstract

Although there has been enormous progress in the last half-century in the drug discovery targeting obesity and associated co-morbidities, the clinical treatment of obesity remains tremendously challenging. GPR75 is an orphan receptor and is suggested to be a potential novel target for the control of obesity and related metabolic disorders. Inhibition of the GPR75 signaling pathway by small molecules, antibodies, or genetic manipulations may provide a therapeutic strategy for obesity. Here, we report the active-like Cryo-EM structure of human GPR75 with an intracellular nanobody, which reveals the receptor activation mechanism. The extensive interaction network required to achieve the active structure helps explain the allosteric coupling between the orthosteric pocket and the G-protein coupling domain. The well-defined orthosteric ligand binding pocket of human GPR75 provides a structural basis for anti-obesity drug discovery.

## Introduction

The prevalence of obesity and associated co-morbidities has become a global healthcare challenge in the 21^st^ century^1^. From the data from WHO, worldwide cases of obesity have nearly tripled since 1975. Obesity is a chronic and degenerative disease associated with other metabolic syndromes and related disorders, like cardiovascular diseases^2^, type 2 diabetes^1^, hypotension^3^, and cancers at a dozen of anatomic sites^4^. Developing anti-obesity medications is tremendously challenging because of multiple adverse side effects observed in the history of clinical treatment^5^. As a result, numerous drugs approved for treating obesity have been withdrawn from the market. Therefore, pharmacological treatment of obesity urgently requires more effective, safer, and long-term medicines to facilitate sustained body weight loss. Although a number of genes that result in severe obesity have been identified^6^, the multiple mechanisms and complex physiological systems of obesity and associated co-morbidities call for new targets and drug development strategies.

GPR75 is a member of the G protein-coupled receptor family and is a novel target for the clinical treatment of obesity^7^. Human GPR75 haploinsufficiency exhibits a striking phenotype of low body fat, and GPR75 knockout mice are hypophagic and thin, improving glucose tolerance and insulin sensitivity^8^. GPR75 was first cloned and identified as an orphan GPCR in the human retinal pigment epithelium and different brain region^9^. A recent result indicates that 20-hydroxyeicosatetraenoic acid (20-HETE), a product of cytochrome P450 (CYP) 4A and 4F isozymes, functions as an endogenous agonist for GPR75^10^. 20-HETE is a potent vasoconstrictor, and upregulation of the production of this compound is related to hypertension and cardiovascular diseases associated with increases in blood pressure^11,12^. A transgenic mouse model overexpressing 20-HETE synthase, together with high-fat diet feeding, displayed hyperglycemia and impaired glucose metabolism^13^. Knockdown GPR75 in a mouse model with 20-HETE dependent hypertension prevented smooth muscle contractility, vascular remodeling, and blood pressure elevation^10^. Meanwhile, blockade of the 20-HETE/GPR75 signaling pathway with 20-HETE mimics lowers blood pressure and alters vascular function in mice^14^. Besides, some studies suggested that the chemokine CCL5 function as an agonist of GPR75^15,16^, playing a role in insulin secretion^17,18^. Collectively, inhibition of GPR75 may provide a therapeutic strategy for obesity and co-morbidities.

Here, we report the Cryo-EM structure of human GPR75 with an intracellular G protein mimic nanobody NbH3 at 3.6 Å. The structural analysis of GPR75 indicated that the receptor is stabilized in an active-like state by the NbH3. The overall structure features of GPR75 are similar to previously reported Class A GPCR, like β_2_AR. However, some famous conserved motifs in Class A GPCR are not conserved in GPR75, which indicates a special conformational allosteric modulation mechanism. In addition, the orthosteric ligand binding pocket of GPR75 is formed by many polar and hydrophobic residues, which may improve the development of in-silico drug discovery.

## Results

### The overall structure of GPR75-NbH3

Due to the heterogeneity of GPCR structure, a specific fab fragment or nanobody has been used to stabilize the GPCR conformation for structural study as previously reported (Fig. S1). We develop a GPR75-specific nanobody by yeast surface display system and evaluate the nanobody binding ability by size exclusion chromatography and 2D classification. We identified a nanobody that specifically binds to the intracellular region of GPR75, as illustrated in the 2D classification result (Fig. S2). A total 14,777 good micrographs were selected for further data processing and were reconstituted to an overall 3.6 Å map (Fig. S2, S3). Fig. 1a shows an overall structure of bril-fused GPR75 in complex with NbH3, which binds on the intracellular surface of GPR75 with the third complementarity-determining region (CDR3) anchoring in the receptor core. A classical orientation of nanobody in the GPCR complex is the CDR3 projects into the core of the receptor nearly vertically ^19–24^, while the NbH3 parallelly floats on the membrane plane. As a result, the CDR3 of NbH3 not only occupies the classical downstream transducer binding pockets, a hotpot epitope like other GPCR-nanobody complexes (Fig. 1b), but also swings into a second cleft formed by ICL1, TM7, and H8. Additionally, several hydrophobic/aromatic residues, including ^Nb^Y103, ^Nb^Y105, ^Nb^L106, and ^Nb^W107, contribute to stabilizing the receptor conformation (Fig. 1c).

**Fig. 1.**
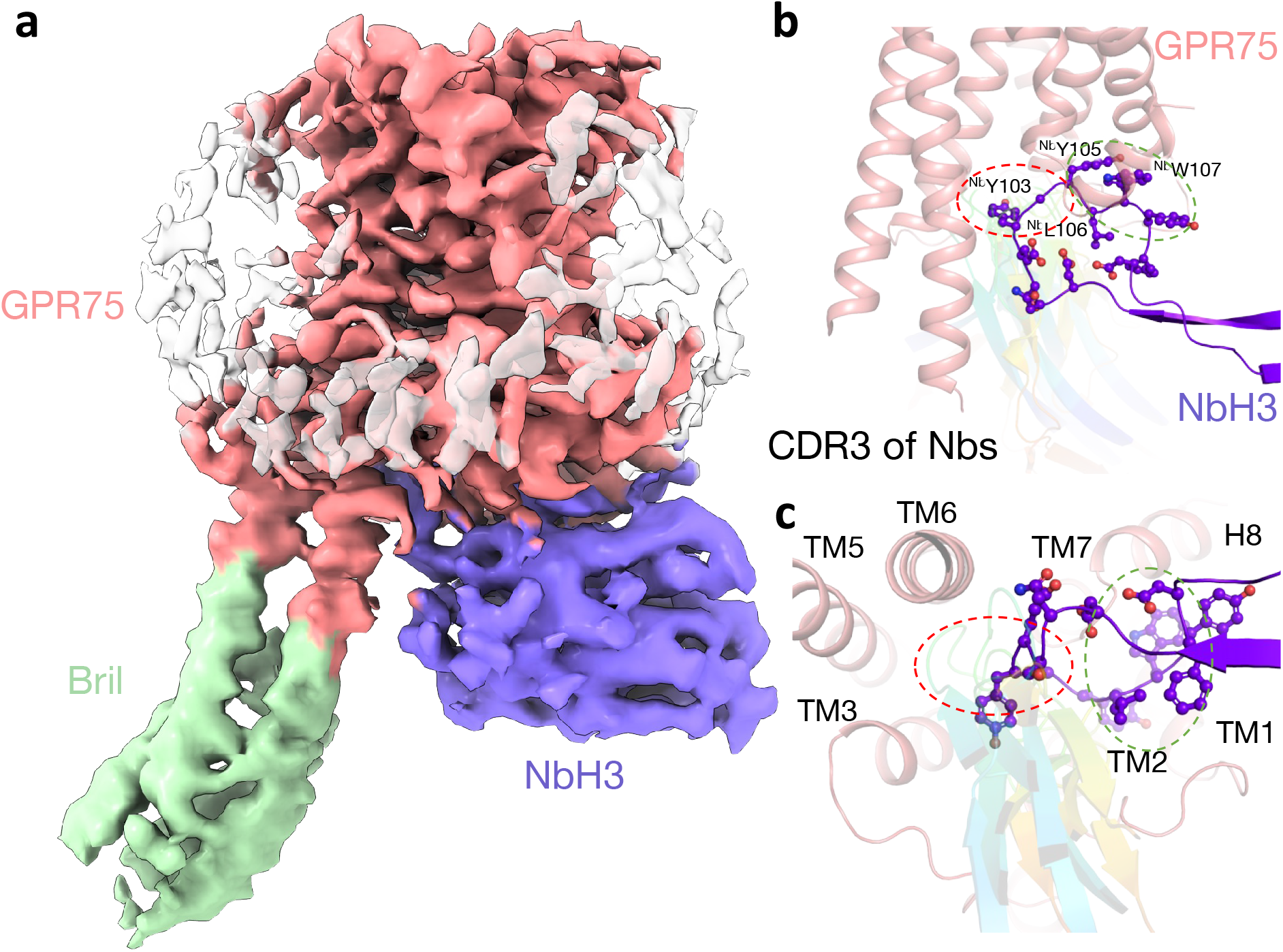
The overall structure of GPR75-NbH3. (a). Orthogonal view of the density map for the GPR75 (salmon) - NbH3 (nitrogen) nanobody complex. The fused bril domain is shown in lime green. (b, c) The CDR3 loop of NbH3 occupied two epitopes, one is the classical epitope shared by a number of GPCR nanobodies (indicated as red dash circle), and the other is a unique epitope formed by ICL1, TM7, and H8 (indicated as green dash circle). The CDR3 of several GPCR-specific nanobodies (PDB ID: 6O3C, 6OS2, 6MXT, 5WB1, 4MQT, 5JQH, 3P0G, 3VG9) is shown as a rainbow cartoon, and residues in NbH3 mediating directly interaction are shown as sticks and spheres.

To interpret the conformational state of the GPR75-NbH3 complex, we superpose the GPR75 structure with the inactive, partially active, and fully active β_2_AR structures, stabilizing by conformational selective nanobodies Nb60^23^, Nb71^25^, and Nb80^24^, respectively (Fig. 2a). To our surprise, the root mean square deviation (r.m.s.d.) over all the transmembrane helices of the β_2_AR (274 Cα atoms) is 4.43 Å, 3.55 Å, and 3.12 Å, respectively. The smaller r.m.s.d. indicates GPR75 is an active-like state. Because we didn’t include ligand in the receptor purification, nanobody screening, and the following Cryo-EM sample preparation step due to the good monomeric behavior in the receptor, the initial expectation is to get the inactive GPR75 structure. For GPCR targets with functional versatility and conformational plasticity, the apo receptor usually prefers to stay in the inactive state according to the energy landscape theory^26–28^. It is unlikely that the active-like conformation of GPR75 is due to bril-fusion because the bril-fused GPCR structures exhibit an inactive structure in the presence of antagonist^29^, and intermediate state^30^ or active state^31^ in the presence of an agonist.

**Fig. 2.**
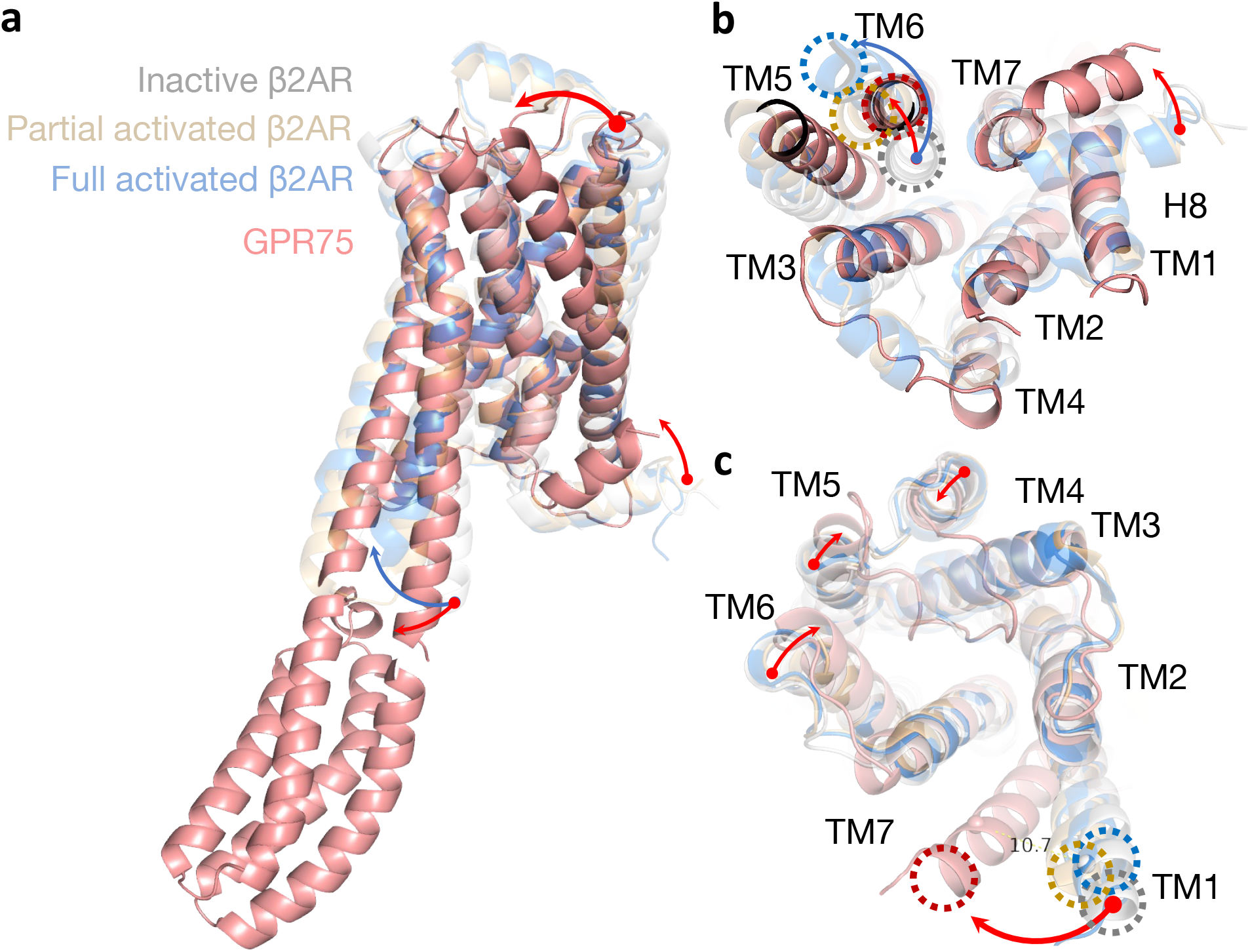
The superposed structures of the GPR75 and three different β_2_AR structures. (a). The overall structure of the active-like GPR75 (salmon) compared with the inactive β_2_AR (grey, PDB ID: 6MXT), partially activated β_2_AR (yellow-orange, PDB ID: 5JQH), and fully activated β_2_AR (light blue, PDB ID: 3P0G). The TM movement of the GPR75 relative to inactive β_2_AR is noted by a red arrow and the TM movement of fully active β_2_AR relative to inactive β_2_AR is noted by a blue arrow.

### The active-like feature of the GPR75 structure

The structural features of GPR75, for example, an outward movement of TM6 compared with inactive β_2_AR, are associated with an active-like state. The TM6 movement is similar to the partially activated salmeterol-β_2_AR-Nb71 complex^25^ (Fig. 2b). Because not all activated GPCR structures present large-scale rearrangements in the cytoplasmic region of TM6, it’s impossible to interpret the activation extent simply by TM6 movement. The subtle inward movement of TM7 and close contact between TM5-TM6 may also reflect a conserved contact rearrangement upon Class A receptor activation^32^. Nevertheless, we can’t exclude the possibility that the close contact between TM5 and TM6 extension may result from the bril fusion design in consideration of the substitution of a long ICL3. Primarily, barcodes on TM5-ICL3-TM6 and TM3-ICL2-TM4 collectively contribute to Gα protein selectivity^33^. The GPR75 is mainly reported to be coupled to downstream Gq protein by both 20-HETE and chemokine CCL5^10,15^, while it is also reported to couple to Gi protein by constitutive G protein coupling profile^34^.

Strikingly, the most significant difference between the GPR75 and β_2_AR structures is found at the extracellular end of the TM1, which moves close to TM7 about 10.7 Å at H38^1.28^ position, relative to D29^1.28^ in the inactive β_2_AR (superscripts in this form indicate Ballesteros–Weinstein numbering for conserved GPCR residues) (Fig. 2c). The conformational change at the extracellular end of TM1 may partially attribute to the binding of NbH3, which caused a large movement of the ICL1 loop relative to TM8. The GxxG motif, which packs close to the conserved NPxxY motif, may play the role of a pivot point that wings the TM1 towards TM7 and enables close contact of TM1 and TM7 in the extracellular end. The extracellular end of TM4, TM5, and TM6 show inward movement compared with the inactive β_2_AR structure, which suggests a contraction of the ligand binding pocket due to the allosteric modulation of extracellular nanobody NbH3 (Fig. 2c).

Class A GPCRs shows a set of common structural rearrangement during receptor activation^32,35^. The extracellular ligand-binding pocket and the intracellular effectors coupling regions are allosterically linked by several well-known but structurally and spatially disconnected motifs, like DRY, NPxxY, PIF, CWxP, and sodium binding pocket. The highly conserved triplet on TM3, the D(E)^3.49^-R^3.50^-Y^3.51^ motif, usually plays a role in maintaining the receptor in an inactive state by forming an intrahelical salt bridge between the R3.50 and E^6.30^ in TM6^36^. For GPR75, the DRY motif, residues H142^3.49^-R143^3.50^-L144^3.51^, is unique and non-canonical in Class A GPCR and is similar to the HRM motif in GPR162 and GPR153^37^. The ionic lock pair of R143^3.50^ and D316^6.30^ is conserved as other GPCRs, and the R143^3.50^ adopts an extended conformation virtually identical to that seen in the β_2_AR-Gs complex^38^ (Fig. 3a). Meanwhile, the D316^6.30^ is far away from the R143^3.50^, which reflects the release of potential structural restraints from TM3. Interestingly, the R143^3.50^ form a cation-π interaction with ^Nb^Y103 as the contact of R131^3.50^ and ^Gs^Y391 in the β_2_AR-Gs complex, which indicates the NbH3 stabilizes the GPR75 conformation in a G protein mimic way (Fig. 3a)^39^. Similarly, the Y376^7.53^ in NPxxY motif in TM7 shows a similar residue arrangement as activated β_2_AR, rhodopsin^40^, and M2 muscarinic receptor^22^, in which a direct or water-mediated interaction between the Y^5.58^ and Y^7.53^ contributes to receptor activation.

**Fig. 3.**
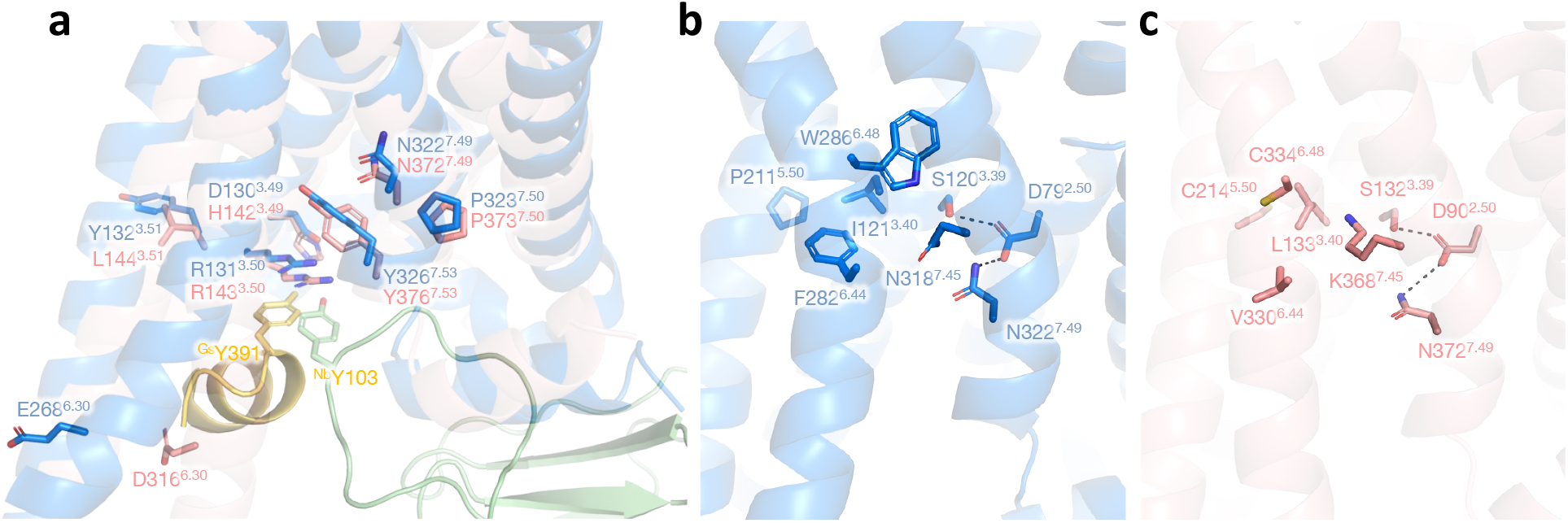
The active-like structural feature of the GPR75. (a). The structural feature of the HRL (DRY) motif and NPxxY motif in the GPR75 (salmon)- NbH3 (green) complex and in the activated β_2_AR (light blue) - Gs (yellow) complex. The ^Nb^Y103 mimics a similar interaction pattern as in the β_2_AR-Gs complex (PDB ID: 3SN6). (b,c). Similar hydrogen bond patterns among TM2, TM3, and TM7 near the CLV (PIF) motif and classical sodium binding sites in the β_2_AR-Gs complex (b, light blue) and the GPR75-NbH3 complex (c, salmon).

The conserved core triad, PIF motif (P^5.50^, I^3.40^, and F^6.44^), located just below the binding pocket (Fig. 3b-c), plays a role in initiating the cascade of structural changes upon receptor activation. One striking feature of GPR75 is the absence of a highly conserved P^5.50^, whose insertion causes a local unwinding of TM5. The C214^5.50^ in GPR75 is non-conservative and only accounts for 1.7% (5 from 292 Homo sapiens GPCRs) in Class A GPCR, the rest four GPCRs are GPR148, LGR5, LGR6, and MRGRE^41^. With the substitution of P^5.50^ by C^5.50^, TM5 shows a more straight and rigid conformation but worse flexibility in response to ligand binding. The alternative version of V330^6.44^ and C334^6.48^ in GPR75, compared with F^6.44^ and W^6.48^ in the β_2_AR, have an irregular small side chain residue, which may confer better allosteric properties and lower the energy barrier for receptor activation.

Another structural rearrangement during Class A GPCR activation is the formation of TM3-TM7 contact^32,42^. In the inactive state of the receptor, a sodium binding site is coordinated by D^2.50^ (92.1% conservative), S^3.39^(71.6% conservative), N^7.45^(65.4% conservative), and N^7.49^(71.9% conservative). The collapse of the sodium binding pocket will lead to a denser repacking of the four residues and initiate the movement of TM7 toward TM3. As shown in Fig. S4, unlike other inactive receptors, there is no space in the classical sodium binding, which also suggests a shrinking interhelical contact. A similar interhelical hydrogen bond network between the GPR75-NbH3 and the β_2_AR-Gs complex implies a structural rearrangement due to the allosteric effect of the intracellular nanobody.

### The ligand binding pocket of the GPR75 structure

The well-known structural plasticity of GPCR is the G protein binding pocket, and the agonist binding pocket is allosterically coupled^43^. Because of the broad diversity of Class A GPCR ligand repertoire, it is not possible to see a common ligand recognizing pattern across all receptors^44,45^. The active-like conformation we observed in GPR75 implies a closed, active, and high-affinity state for agonists. We observed continuous density in the orthosteric ligand binding pocket, but it’s hard to define whether it’s the ICL2 loop density or unknown ligands (Fig. 4a-b). The ICL2 loop possibly forms a lid over the ligand binding pocket to modulate initial ligand recognition^46^. The ICL2 in other Class A GPCR shows highly differentiated structures, forming helices, sheets, or intrinsically disordered loops. Based on the position of the conserved disulfide pair between C118(TM3) and C188(ICL2), we suspect that the density map in the pocket may come from the ICL2 loop, while we can’t extrude the possibility that it is unknown molecule coming from cells or purification conditions. Although, due to the low resolution in the extracellular surface region, we can’t well define a full model of all ECL loops, the residue density of GPR75 in the orthosteric ligand binding pocket is well enough to identify the valuable pocket.

**Fig. 4.**
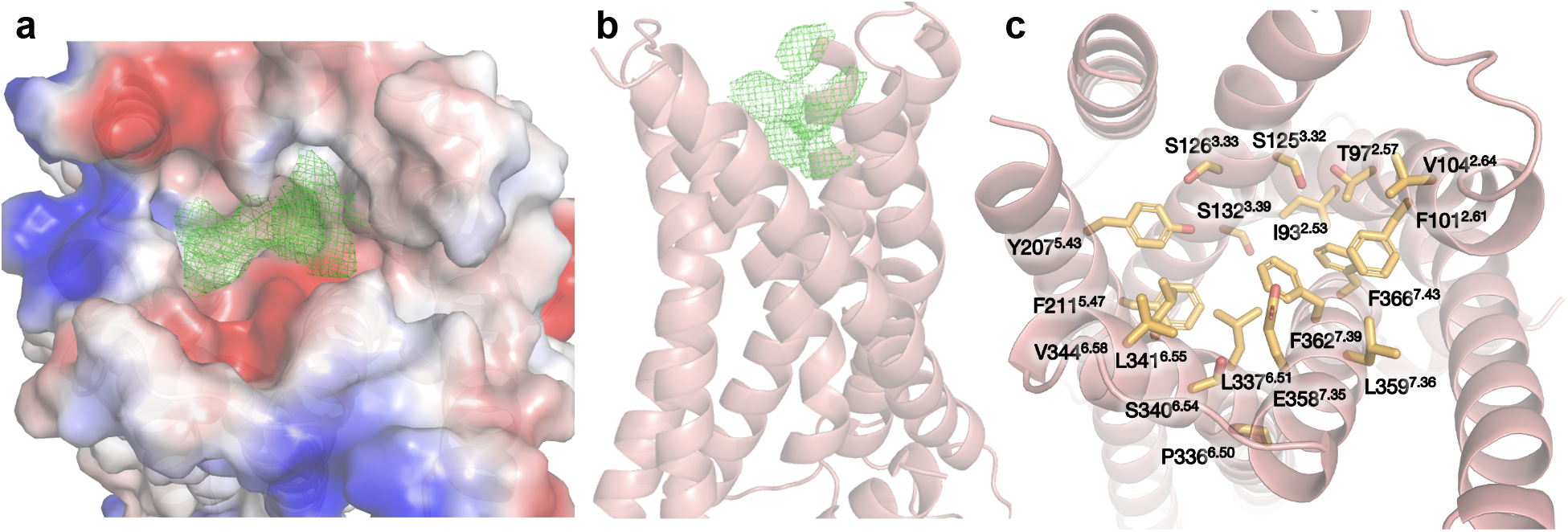
The orthosteric ligand binding pocket of GPR75. (a). Top surface view of the density in the orthosteric ligand binding pockets. (b). Top surface view of the orthosteric ligand binding pockets. (c). Some hydrophobic and polar residues are involved in forming the orthosteric ligand binding pockets.

Compared with the ligand binding pocket of activated β_2_AR and GPR75, the latter show shallow, spacious pockets with slightly negatively charged (Fig. 4a-b, S5). The pocket is on top of the PIF (as C^5.50^/L^3.40^/V^6.44^) and CWxP (C333^6.47^/C334^6.48^/P336^6.50^) motif to facilitate conformational rearrangement upon agonist binding. Consistent with the consensus scaffold interface for ligand binding^35^, the major pocket of GPR75 is formed by the extracellular end of TM2, TM3, TM5, TM6, and TM7, packaged by a number of polar and hydrophobic residues. The polar residue cluster, S125^3.32^, S126^3.33^, S132^3.39^, Y207^5.43^, Y207^5.43^, and E358^7.35^, may contribute to the endogenous ligand interaction in cells in a manner mentioned before^44^ (Fig. 4c). Historically, various 20-HETE-related pharmaceutical agents have been synthesized, including 20-HETE agonists and antagonists^47^. The structure-activity relationship analysis of 20-HETE analogs indicates that 20-HETE agonists and antagonists require a carboxyl or an ionizable group on carbon 1 and a double bond near the 14 or 15 carbon. Meanwhile, 20-HETE agonists also require a functional group capable of hydrogen bonding on carbon 20 or 21, whereas antagonists lack this reactive group^48^. It should be noted that three mutations, S205^5.41^, T212^5.48^, and S219^5.55^, predicted to involve 20-HETE-GPR75 interaction, are not located in the central cavity of orthosteric ligand binding pockets of the active-like structure of GPR75^49^.

### Conclusion

The GPR75 protein-truncating variants in large-scale human populations are genetically associated with lower body mass index^7^. The active-like Cryo-EM structure of GPR75-NbH3 here provides a clue to revealing the receptor activation mechanism, which is critical for developing novel therapeutic anti-obesity drugs. Comparison of active-like GPR75 structure with other active-state Class A GPCR structures offers insights into shared mechanisms for receptor activation. The extensive interaction network required to achieve the active structure helps explain the allosteric coupling between the orthosteric pocket and the G-protein coupling interface. Considering the physicochemical property of the reported agonist, 20-HETE, it looks reasonable that the shallow hydrophobic pockets will fit the ligand and initiate conformational transmission in a manner observed in the other Class A receptors. Although several 20-HETE derivative antagonists have been developed, their fatty acid property with a rather high albumin binding rate in the plasma may restrict the distribution of these compounds to targeted tissues. The structural-based drug design targeting GPR75 may accelerate the discovery of a lead compound with novel properties^50^. Besides, therapeutic approaches based on genetic manipulations, such as siRNA oligonucleotide, also provide an alternative therapeutic intervention for the treatment of obesity and associated co-morbidities by targeting the GPR75.

## Method

### Expression and purification of the human GPR75 receptor

A truncated human GPR75 receptor (residue1-395) bearing an amino-terminal haemagglutinin signal sequence followed by the FLAG epitope, and a carboxy-terminal Strep-tag and 6xHis tag was cloned into pfastbac-1 vector. To enhance the expression level of GPR75, twenty-four amino acids from the β_2_AR receptor (MGQPGNGSAFLLAPNRSHAPDHDV) were used to replace the original residues1-31 in a modified version of the GPR75 receptor. Furthermore, the bril sequence was used to replace the original ICL3 loop (237-306) to enhance receptor stability. The receptor was expressed in *Sf9* insect cells using the Bac-to-Bac baculovirus system. The expression and purification of the GPR75 receptor are according to the methods described previously.^38^ Briefly, the GPR75 receptor was expressed for 48 hours after infection with recombinant baculovirus. Cells were lysed, and extracted using a buffer containing 20 mM Tris, pH 7.5, 750 mM NaCl, 0.5% lauryl maltose neopentyl glycol (LMNG), 0.03% CHS, 0.2% sodium cholate, 2.5 μg/ml leupeptin. Ni-NTA affinity purification was used as the initial purification step and followed by Flag affinity chromatography for further purification. The eluted receptors were loaded onto a Superdex 200 column equilibrated in a buffer containing 20 mM Tris, PH 7.5, 150 mM NaCl, 0.002% LMNG, 0.00015% CHS. Subsequently, the concentrated sample was then aliquoted, flash-frozen, and stored at −80°C.

### Isolation of nanobody binders from library

Nanobodies were selected from a synthetic yeast display nanobody library^51^. In the first round of screening, 5×10^9^ cells of the yeast display nanobody library were washed and resuspended in 2 mL selection buffer (20 mM Tris pH7.5, 150 mM NaCl, 0.06% LMNG, 0.003% CHS, 0.02% sodium cholate, 5 mM MgCl_2_) and then incubated with FITC-labeled anti-Flag M1 antibody and anti-FITC microbeads (Miltenyi) at 4°C for 40 min. Non-specific binding nanobodies were removed by pre-clear which involved passing the yeast through an LD column (Miltenyi), and the remaining yeast from the flow-through was incubated with 400 nM GPR75 for 30 min at 4°C, and then washed once with 2 mL selection buffer. Yeast was stained with 200nM FITC-labeled anti-Flag M1 antibody at 4°C for 20 min, followed by washing with selection buffer to remove excess M1 antibody. After incubation with anti-FITC microbeads at 4°C for 20 min, nanobodies specifically bound to GPR75 were enriched by passing through LS column (Miltenyi), and cultured for 24 hours in -TRP medium at 30°C. Rounds 2 of MACS selection were performed similarly with 1×10^8^ yeast, and cells were washed and resuspended in 500μL buffer, incubated with 200 nM GPR75, stained with FITC-labeled anti-Flag antibody M1 and anti-FITC microbeads.

After 2 rounds of MACS, the diversity of yeast display nanobody library was less than 10^6^. In order to test the enrichment effect, 10^6^ cells were stained with 100 nM GPR75, 100 nM FITC-labeled anti-Flag antibody M1, and Alexa Fluor 647-labeled anti-HA antibody, ~3.3% of the MACS2 pool was positive for GPR75 binding. The enriched yeasts were used for further selection by FACS. 10^7^ cells were incubated with 200 nM GPR75 in 100μL selection buffer at 4°C for 1h. After incubation, yeast cells were washed twice with ice-cold selection buffer, then incubated with 100 nM FITC-labeled anti-Flag antibody M1 and 0.5 μg Alexa Fluor 647-labeled anti-HA antibody (Cell Signaling Technology) in 100μL selection buffer at 4°C for 20 min. After incubation, yeast cells were washed three times with ice-cold selection buffer, suspended in 1mL of selection buffer, and sorted on FACSAria (BD). Typically, 0.5% of the GPR75 binding population was gated for collection. Collected cells were grown in -TRP medium, and about 15% of the FACS1 pool was positive for GPR75 binding. After FACS2, plate 2×10^4^ cells in -TRP agar and separate for a single colony for sequencing.

### Expression and purification of nanobody and GPR75-nanobody complex

The isolated nanobody sequence was cloned into the pET26b vector with an amino-terminal PelB leader sequence (MKYLLPTAAAGLLLLAAQPA) for periplasmic protein expression and with a C-terminal 6xHis-tag, and transformed into *E. coli* cells BL21(DE3). Cells were induced in Terrific Broth medium with 1 mM IPTG at OD600 of 1.2 and cultured with shaking at 22°C for 20 h. Periplasmic protein was obtained by osmotic shock, and the nanobodies were purified using Ni-NTA chromatography, followed by a Superdex 200 column equilibrated in buffer (20 mM HEPES, pH 7.5, 150 mM NaCl). The eluted sample was concentrated, aliquoted, flash-frozen, and stored at −80°C.

### Preparation of GPR75-nanobody complex and EM data acquisition

The complex was formed by mixing the receptor with 5x excess of the selected nanobody in a buffer condition (20mM Tris, PH 7.5, 150mM NaCl, 0.002% LMNG, 0.00015%CHS). The complex was preincubated for 1 hour on ice before loading to a pre-equilibrated Superdex 200 column. The eluted sample was concentrated for grid preparation.

An aliquot of 4 μL protein sample of GPR75-NbH3 complex at a concentration of 7.6 mg/ml was applied onto a glow-discharged 300 mesh grid (Quantifoil Au R1.2/1.3), blotted with filter paper for 3.0 s and plunge-frozen in liquid ethane using a Thermo Fisher Vitrobot Mark IV. Cryo-EM micrographs were collected on a 300kV Thermo Fisher Titan Krios G3i electron microscope equipped with a K3 direct detection camera and a BioQuantum image filter (GIF: a slit width of 20eV). The micrographs were collected at a calibrated magnification of x130,000, yielding a pixel size of 0.27 Å at a super-resolution mode. In total, 14,777 micrographs were collected at an accumulated electron dose of 50e^-^Å^-2^s^-1^ on each micrograph that was fractionated into a stack of 32 frames with a defocus range of −1.0 μm to −2.0 μm.

### Cryo-EM data processing, model building, and refinement

Beam-induced motion correction was performed on the stack of frames using MotionCorr2^52^. The contrast transfer function (CTF) parameters were determined by CTFFIND4^53^. A total 14, 777 good micrographs were selected for further data processing using cryoSPARC^54^. Particles were auto-picked by the blob picker and template picker program in cryoSPARC, followed by 3 rounds of reference-free 2D classifications. Next, 1, 338, 445 particles were selected from good 2D classes and were subjected to 3 rounds of muti-reference 3D classification using starting models generated using conventional 3D classifications. One converged 3D class from each round of muti-reference 3D classifications with a feature containing GPR75-Bril-NbH3 was selected and removed duplicates. A last heterogeneous refinement was performed and 503,557 particles from a 3D class showing the highest resolution feature were selected for a round of 3D refinement, yielding a final reconstruction at a global resolution of 3.64 Å based on the gold-standard Fourier shell correlation criterion at FSC=0.143. The local resolution was then calculated on the final density map.

The model of the GPR75-NbH3 complex was built by fitting a structure of the complex (predicted by AlphaFold2^55^ and CryoNet) into the density map using UCSF Chimera^56,57^, followed by a manual model building of the complex molecules in COOT^58^ and a real space refinement in PHENIX^59^. The model statistics were listed in Supplementary Table 1.

## Acknowledgments

We thank Pan Li and Haibin Liu from Shuimu BioSciences for their guidance in the model building with CryoNet. We thank Andrew Kruse from Harvard Medical School for kindly providing the synthetic yeast nanobody library.

## Contributions

Z.L., Y.H., H.G., and D.H. performed experiments, Z.L., S.Z., B.L. performed nanobody screening experiments, X.L. supervised nanobody screening, Y.X., F.M., Y.W., and H.Z. prepared grid, collected data and processed data, J.L. built the model and assisted structure analysis, J.L., W.Z., Y.L., and J.H. supervised experiments and analyzed the data, X.N. supervised Cryo-EM data processing, J.H. wrote the manuscript.

## Supplementary Figures

**Fig S1.**
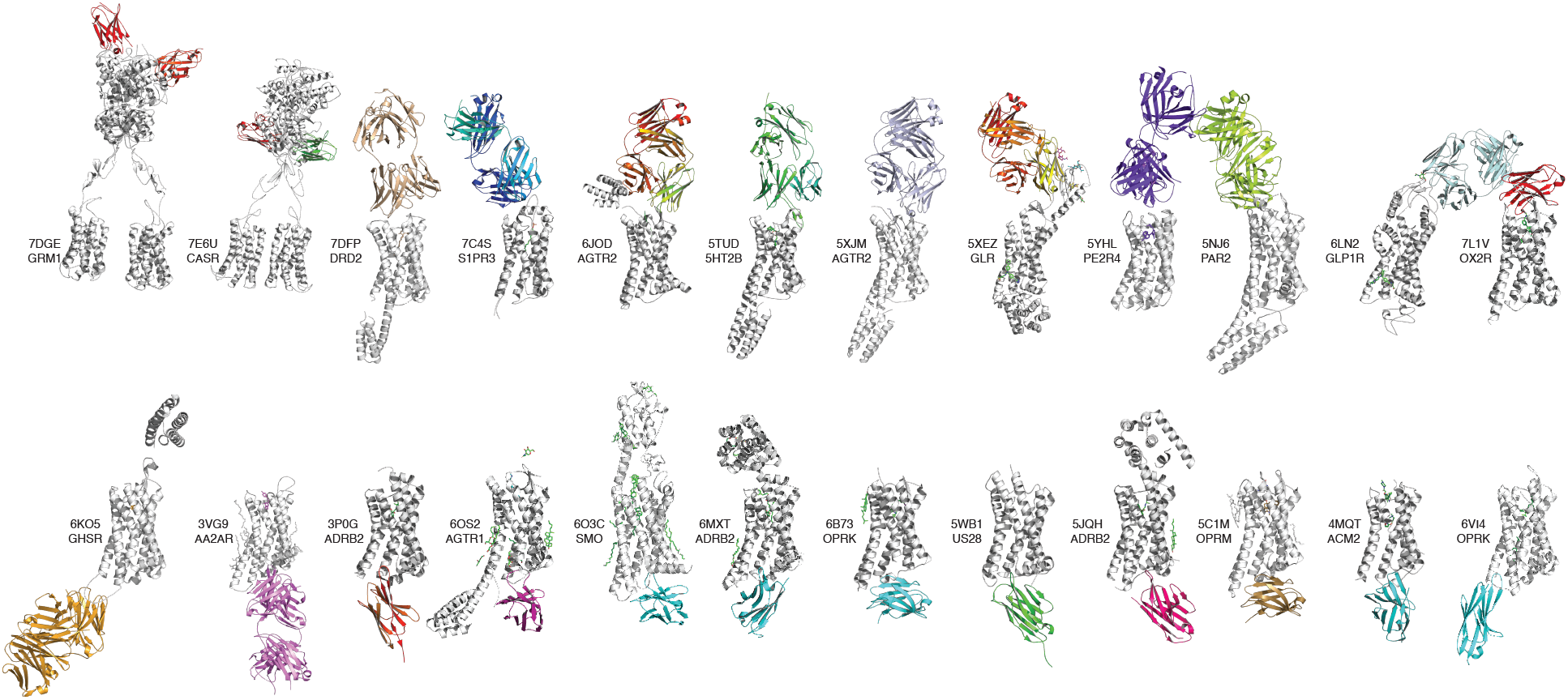
Collective structures of GPCR and fab fragment or nanobody. Receptors are shown as grey cartoons and the fab fragment or nanobody as colored cartoons.

**Fig S2.**
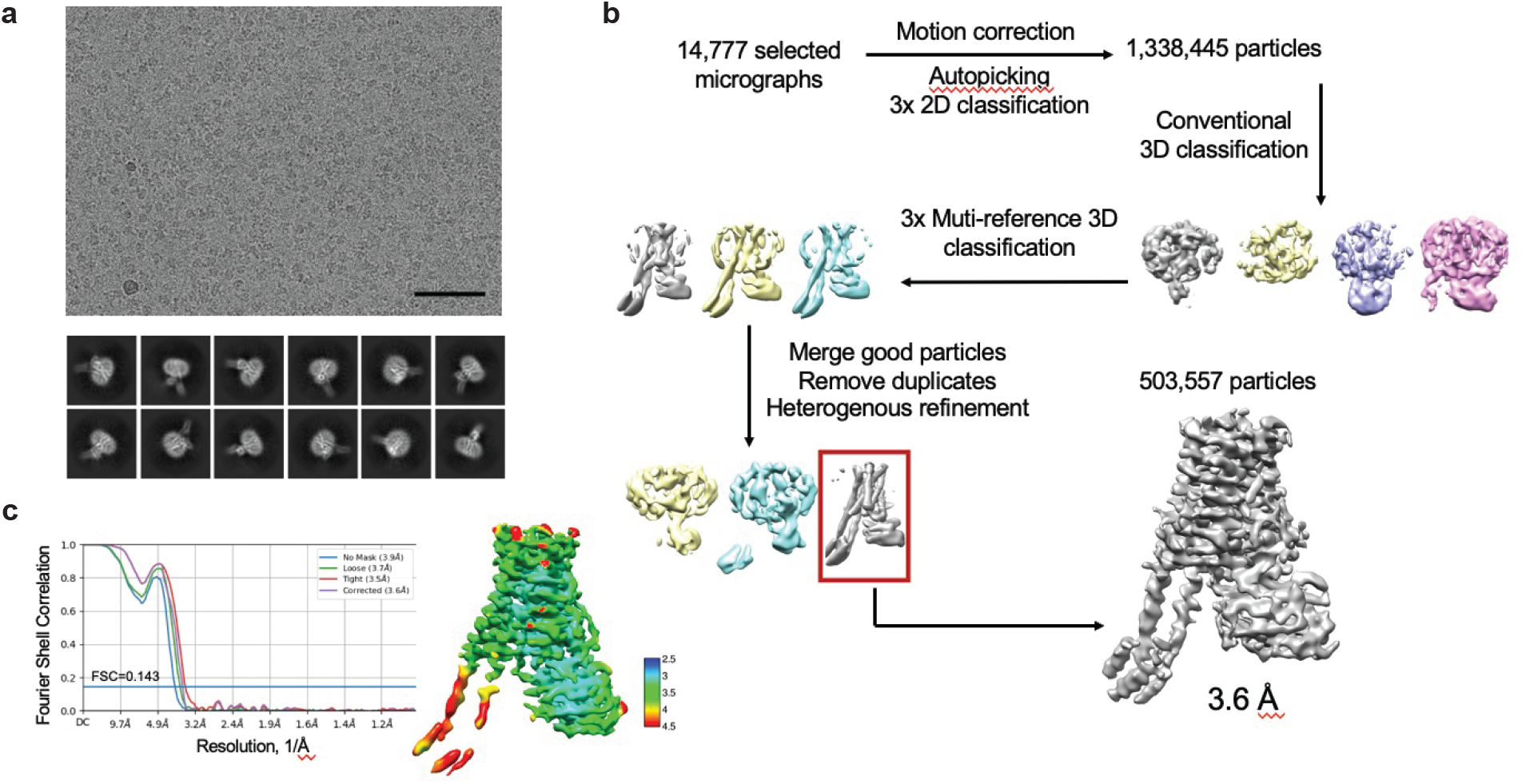
Cryo-EM analysis of the GPR75 complex. (a). Representative electron micrograph and 2D class averages. The black scale bar in the top panel represents 50nm. (b). Flowchart for EM data processing. Details can be found in Methods. (c). The gold-standard Fourier shell correlation (FSC) curve for the final 3D reconstruction (left panel); Local-resolution map for the 3D EM reconstruction of GPR75 complex (right panel).

**Fig S3.**
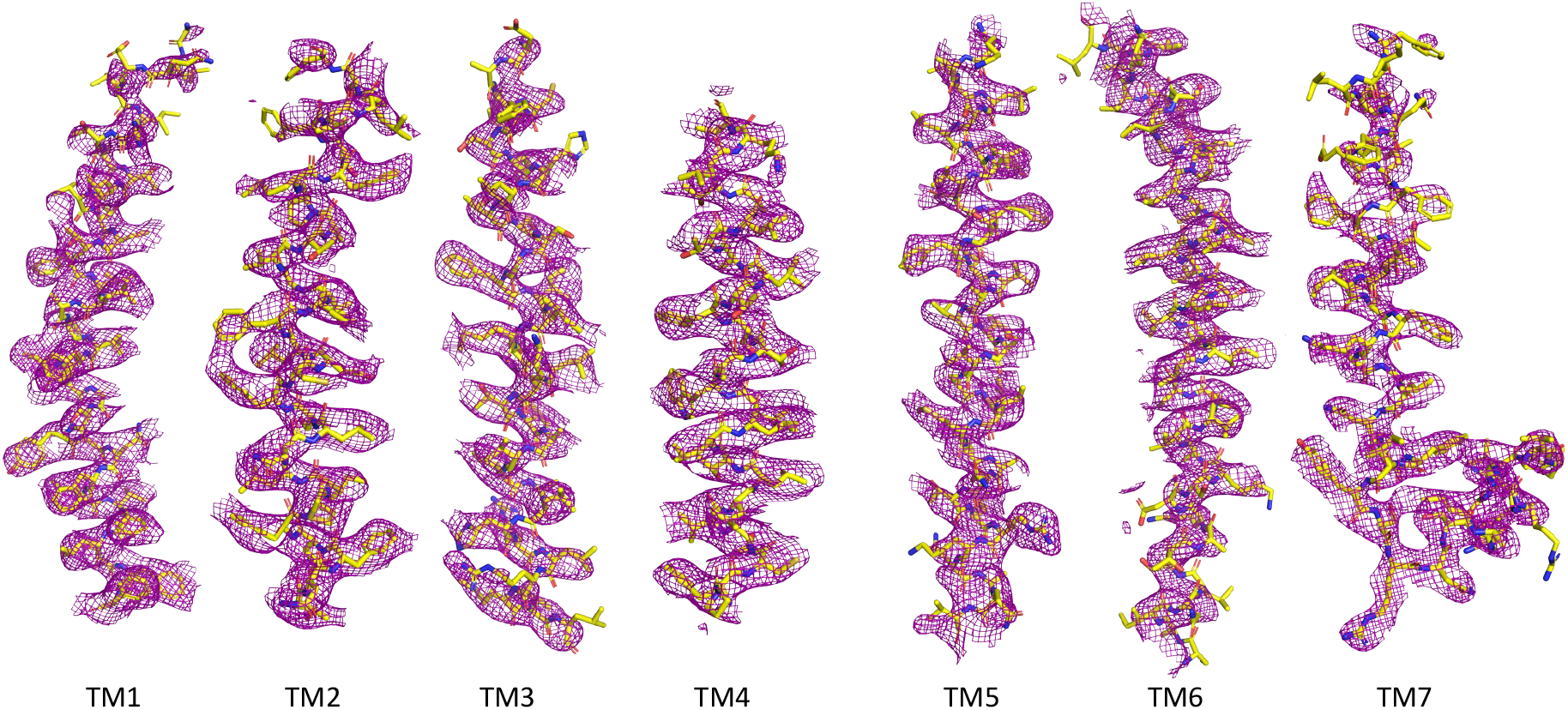
Representative cryo-EM densities of seven transmembrane helices. Representative seven transmembrane helices regions of the GPR75 model are shown as yellow cartoons and the density from the electron microscopy map as red mesh.

**Figure S4.**
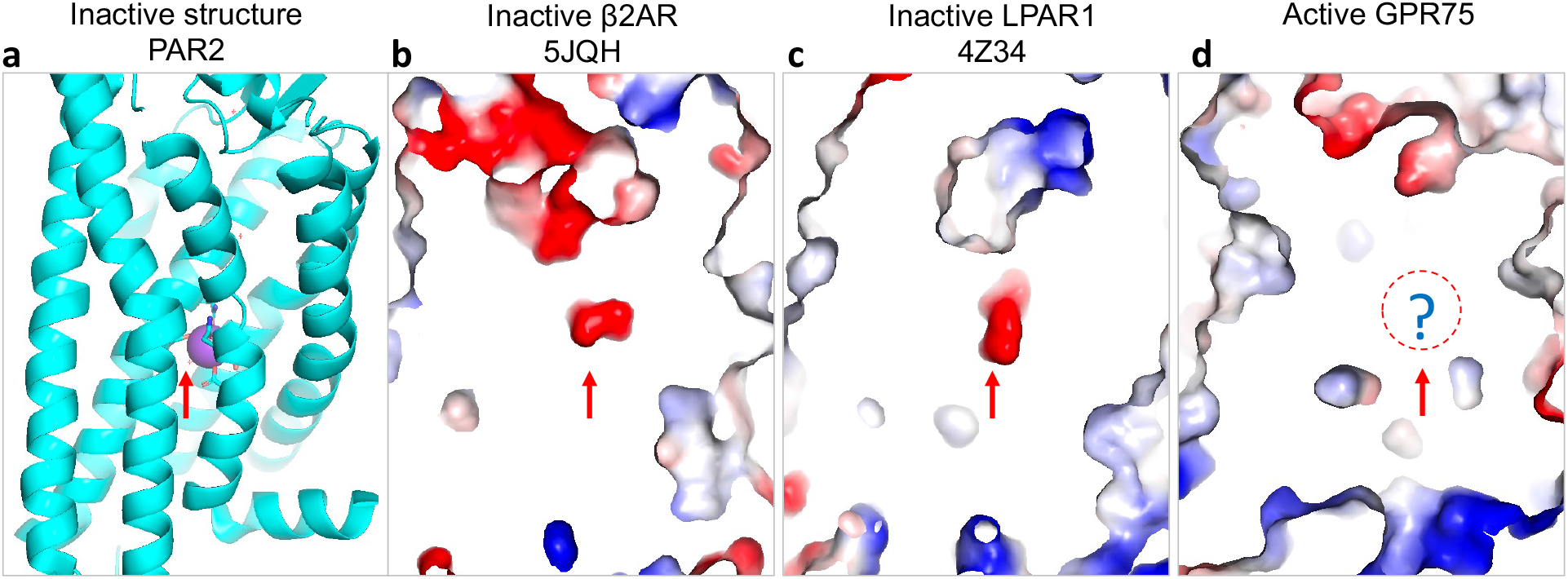
Shrinking sodium binding pocket in the active-like state of GPR75. In the representative inactive Class A GPCR structures, there is a space for sodium binding, while for the active GPR75, the sodium binding site may shrink due to the rearrangement.

**Figure S5.**
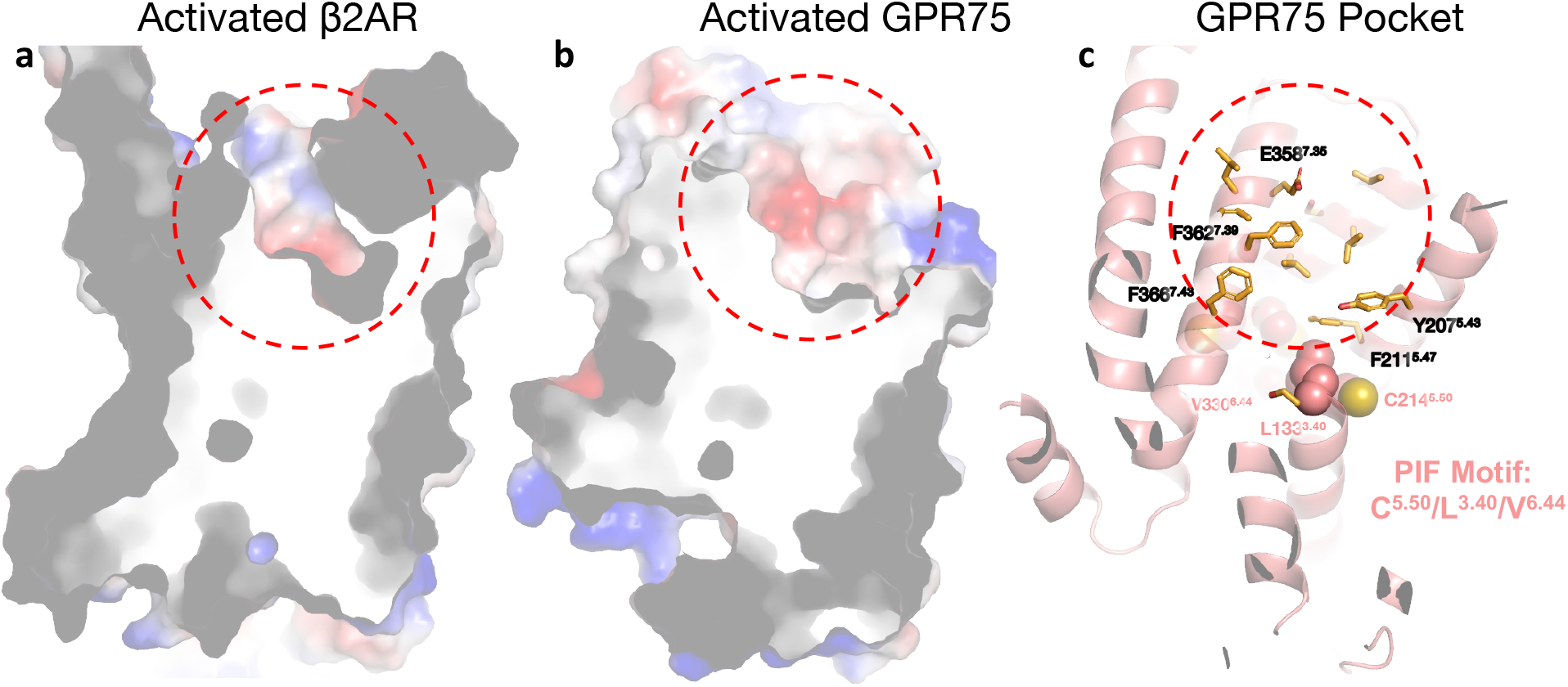
The ligand binding pocket of GPR75. (a-b). The activated β_2_AR shows a narrow ligand binding pocket, while the GPR75 exhibits a large and shallow pocket. (c). The pocket is located near the C214^5.50^ and is formed by a number of hydrophobic residues. The PIF motif is noted as salmon spheres.

**Supplementary Table 1.** Cryo-EM data collection, refinement, and validation statistics

|  | GPR75-NbH3 Consensus map |
| --- | --- |
| **Data collection and processing** |  |
| Magnification | 130k |
| Voltage (kV) | 300 |
| Electron exposure (e<sup>-</sup>/Å<sup>2</sup>) | 50 |
| Defocus range (μm) | -1.0~-2.0 |
| Pixel size (Å) | 0.54 |
| Symmetry imposed | C1 |
| Initial particle images (no.) | 2,968,809 |
| Final particle images (no.) | 503,557 |
| Map resolution (Å) | 3.64 |
| FSC threshold | 0.143 |
| **Refinement** |  |
| Initial model used (PDB code) | AF2 predicted |
| Model resolution (Å) | 3.60 |
| FSC threshold | 0.143 |
| Map sharpening *B* factor (Å<sup>2</sup>) | -230 |
| Model composition |  |
| Protein residues | 523 |
| R.m.s. deviations |  |
| Bond lengths (Å) | 0.003 |
| Bond angles (°) | 0.678 |
| Validation |  |
| Clashscore | 10.90 |
| Poor rotamers (%) | 0.23 |
| Ramachandran plot |  |
| Favored (%) | 97.11 |
| Allowed (%) | 2.70 |
| Disallowed (%) | 0.19 |

